# Unveiling the phenology and associated floral regulatory pathways of *Humulus lupulus* L. in subtropical conditions

**DOI:** 10.1101/2023.09.15.557988

**Authors:** Robert Márquez Gutiérrez, Raphael Ricon de Oliveira, Thales Henrique Cherubino Ribeiro, Kellen Kauanne Pimenta de Oliveira, João Victor Nunes Silva, Tamires Caixeta Alves, Laurence Rodrigues do Amaral, Marcos de Souza Gomes, Matheus de Souza Gomes, Antonio Chalfun-Junior

## Abstract

The *Humulus lupulus* L. (hops) is traditionally produced in temperate regions, historically attributed to the belief that vernalization was necessary for reproductive transition. Nevertheless, recent studies have revealed the potential for hop plants to flower in tropical and subtropical climates, thereby reshaping cultivation prospects once the developmental disparities are understood and adapted for optimized production. Here, we delved into the hop phenological cycle and the underlying floral regulatory pathways ultimately, seeking to devise strategies to modulate development and enhance production in regions with low latitudes. In contrast to cultivation in temperate regions, hops grown in the subtropical climate of Minas Gerais, Brazil, flower multiple times throughout the year, independently of the season. Hop is a short-day plant, and in Brazil, the photoperiod is consistently inductive due to daylight hours always being below the described threshold (16,5 h of light critical point). We observed that the reproductive transition of hop initiates after a specific number of nodes, 25 to 28, leading us to hypothesize that this process is primarily controlled by endogenous factors. To explore this issue, we first identified in the hop genome families of miRNAs related to reproductive transition, such as the families MIR156 and MIR172, and the PEBP family which includes the FLOWERING LOCUS T (FT). The expression of miR156 decreased along development inversely to miR172 and in synchrony with the upregulation of HlFT3 and HlFT5. These findings strongly support that the reproductive transition in hops under inductive photoperiodic conditions is primarily regulated by endogenous factors related to development and aging throughout the miRNA and FT-associated pathways. Thus, our study shedded light on the intricate molecular mechanisms that underlie hop floral development paving the way for potential advancements in hop production on a global scale.

## 1. Introduction

*Humulus lupulus* L. (hops) is an herbaceous, dioecious and perennial liana belonging to the Cannabaceae family (Natsume et al., 2015). The main product of this plant is the female inflorescences (cones or strobila) that present lupulin producing glands, resin rich in aromatic compounds and essential oils used in the manufacture of various products, including beer (Verzele and Keukeleire, 1991). The α and β acids present in the resins provide bitterness and aroma to the beer. Despite its economic importance, hops have been mostly cultivated in temperate zones between latitudes 35° and 55° N or S of the Equator (Jastrombek et al., 2022) and few works have explored its phenological cycle in association with the regulatory networks governing flowering in tropical and subtropical regions.

The hop phenological cycle includes the establishment of rhizomes until the production of flowers and senescence, a period correlated with variable environmental conditions in temperate regions (Roßbauer et al., 1995). The hop rhizome is perennial, with a longevity of approximately 14 years, and the aerial part (stem, leaves and flowers) is annual (Fric et al., 1991). Hop growth in temperate regions begins after 6 months of inactivity (winter) at the time corresponding to early to mid-spring. Vegetative growth continues in late spring and throughout the summer. And when the days begin to shorten in late summer and early autumn, vegetative growth ceases and the first reproductive structures begin to develop, which is followed by very rapid growth of the cones during at least a month. By early autumn, the plants translocate all nutrients to the roots and the aerial part perish (Roßbauer et al., 1995).

Hop is a plant that grows optimally in photoperiod above 13 light hours (h), is a short-day plant (16.5 h critical), and the flowering is achieved once the plants reach a determined size (Thomas and Schwabe, 1969). Recently, it has been demonstrated that hop plants do not require vernalization to floral transition and rhizome age does not influence hop yields (Bauerle, 2019). In agreement, no members of the FLOWERING LOCUS C (FLC) orthologs, a known flowering repressor related to vernalization (Zhu et al., 2015), were found in the hop genome (Márquez Gutiérrez et al., 2022).

The phenology of four hop cultivars grown in Santa Catarina (a state in the south of Brazil) was evaluated, being described that the vegetative growth occurs between September and December while the reproductive stage takes place at the end of December and the cones development between January and February of the subsequent year (Fagherazzi et al., 2023). Some of the phenology stages proposed are not found in subtropical regions, for example, hop plants are capable of blooming prematurely, skipping the lateral shoot formation, stages that are a normal step in the development in temperate latitudes. This causes unevenness in the cones and therefore impairs the harvesting in the sub-tropics (Acosta et al., 2021).

The dynamic of some microRNAs (miRNAs) transcription is related to floral transition (Schwab, 2012). miRNAs are a type of small non-coding RNA (generally with 20-22 nt in length) that post-transcriptionally regulates the expression of mRNAs target genes through RNA–RNA interactions (Teotia and Tang, 2015). During the juvenile phase, miR156 accumulates in leaves of *Arabidopsis thaliana* to repress transcription factors of the SQUAMOSA PROMOTER-BINDING PROTEIN-LIKE family (SPL) (Wang, 2014; Wu et al., 2009). In some plants, the transition of vegetative to reproductive phase is accompanied by leaf morphology changes, known as heteroblasty, e.g, *Pisum sativum* (Vander Schoor et al., 2022), *Passiflora edulis* (Silva et al., 2019), *A. thaliana* (Wu et al., 2009) which is also observed in hop (Fric et al., 1991). The identity of leaves is controlled by SPL transcription factors expressed in the shoot apical meristem (SAM) during the early vegetative development (Fouracre and Poethig, 2019). Recently it was demonstrated that the mechanism causing the miR156 down-regulation is due to epigenetic marks added in the MIR156 locus elicited by cell divisions in the SAM (Cheng et al., 2021). Therefore, the accumulation of miR156 in leaves, which originated from the SAM, works as a timer that reflects the plant’s aging (number of cells division).

On the other hand, acting antagonistically to miR156, there is the miR172 that accumulates during leaf development and promotes floral transition (Schwab, 2012; Teotia and Tang, 2015). The miR172 is up-regulated by SPLs and, in its turn, represses the APETALA2 (AP2), a transcriptional repressor of *FLOWERING LOCUS T* (*FT*) (Aukerman and Sakai, 2003). Accordingly, in Arabidopsis the expression of *FT* is inhibited by AP2 or AP2-like transcription factors (Castillejo and Pelaz, 2008; Hu et al., 2021; Mathieu et al., 2009) in a positive feedback that also regulates the miR156 expression (Yant et al., 2010). Therefore, *FT* is responsive to diverse endogenous and environmental stimuli perceived by molecular pathways in leaves before the FT protein can be translocated to the meristems to induce flowering (Corbesier et al., 2007).

Orthologs of FT have been identified in several species such as rice (Tamaki et al., 2007), coffee (Cardon et al., 2022) and many others (Pin and Nilsson, 2012). FT belongs to the PHOSPHATIDYLETHANOLAMINE-BINDING PROTEIN (PEBP) family, which is conserved in plants and animals (Jin et al., 2021). In plants this family is divided into four subfamilies: Mother of Flowering Locus T (MFT-clade), Brother of Flowering Locus T (BFT-clade), Flowering Locus T (FT-clade) and Terminal Flower (TFL-clade) (Jin et al., 2021). The presence of specific amino acids in the FT protein is critical to induce flowering (Ahn et al., 2006; Ho and Weigel, 2014). In *A. thaliana*, the expression of *FT* in leaves is promoted by long day conditions, enhanced by the presence of multiple cis-regulatory elements (CREs) occurring at least 7000 nt upstream of the FT locus (Adrian et al., 2010). For example, one of these CREs is the E-box domain which enhances the *FT* expression in response to blue light (Liu et al., 2008).

The molecular network complex, represented by miR156/172 and *FT*, was found to be conserved in multiple plant species such as *P. edulis* (Silva et al., 2019), *Dendrobium catenatum* (Zheng et al., 2019) and *P. sativum* (Vander Schoor et al., 2022). Thus, the interplay between age, photoperiod and other molecular pathways influences the miR156-SPLs and miR172-AP2 targeting pairs levels and, under an optimum balance, promotes the *FT* expression and, consequently, the reproductive transition (Teotia and Tang, 2015). Despite its importance for floral induction, this regulatory module was not yet explored in hop.

In this work, we first characterized the phenological stages of hop development under subtropical conditions of Brazil. Then, we identified conserved MIRNA families and members of the PEBP family, with special focus on *FT-like* genes, in the hops cascade genome (Padgitt-Cobb et al., 2021). Finally, we analyzed the expression profile of miR156, two miR172 and two *FT-like* throughout the hop development. Since the miR156-172 and FT-like constitute a floral regulatory network, our results demonstrate that hop flowering is mainly controlled by endogenous factors related to age and development when cultivated in photoperiod inductive conditions (less than 16.5 h of light). Thus, this work contributed to the better understanding of *H. lupulus* L. reproductive development in the tropics, opening perspectives to improve hops production worldwide.

## 2. Materials and methods

### 2.1. Plant material and field conditions

To determine the phenological stages and the transcriptional profile of miR156 and miR172 in subtropical conditions, female plants of three hop cultivars were used: Chinook, Millenium and Northern Brewer (NB). The experiment was conducted in one-year-old plants (during the years 2019-2021) in a locality of Lavras/MG-Brazil (−21.246732, −45.002251; 918 m.s.n.m). Lavras is a region considered to have a subtropical climate marked by a rainy temperate (mesothermal) with dry winters and rainy summers, and temperature of the hottest month above 22 °C in February (Dantas et al., 2007). The soil used was analyzed by the Soil Analysis Laboratory in the Department of Soil Sciences at Federal University of Lavras (UFLA) and classified as clayey (36 % of sand, 18 % of silt and 46 % of clay), with a content of organic matter of 1.23 %, a cation exchange capacity of 7.20 cmolc/dm3, and pH of 6.4. Each cultivar was planted in rows separated by 1.5 m and each plant 1 m apart. Three stems per plant grew over fiber strings of 0.5 cm of diameter. The crop management was done according to Dodds, (2017). To analyze the dynamics of miR156 and miR172, pairs of leaves of the 5th, 10th, 15th, and 20th nodes were collected from plants with size of 10, 15, 20 and 25 nodes, respectively, immediately immersed in liquid nitrogen and deep freezer (−80° C) stored until RNA extraction.

To evaluate the expression of *FT-like* genes, hop plants of the Northern Brewer (NB) genotype were grown in a Greenhouse of Plant Physiology sector of Federal University of Lavras. The plants were produced according to Bauerle, (2019), firstly the cuttings were immersed in Hoagland solution (1/4) during two weeks in growth chambers (16/8 light/obscure to 22 °C) until the cuttings developed roots. Secondly, the plants were transferred to 11 L pots with a mixture of 2:1 soil-sand and maintained in greenhouse condition (28/20 °C day/night). The plants grew during October to December which the photoperiod extends from 11.40 to 13.5 h light. The foliar material was collected from plants with 10, 15, 20, and 25 nodes during the dusk and stored as was previously mentioned.

### 2.2. miRNAs prediction in hop genome

The *H. lupulus* assembly genome and genome annotation (Padgitt-Cobb et al., 2021) files were obtained from hopbase (http://hopbase.cgrb.oregonstate.edu/). The identification of microRNAs was done according to de Souza Gomes et al. (2011). Sequences identified as precursors of miRNA molecules were structurally and thermodynamically characterized using RNAfold software (ViennaRNA Services, http://rna.tbi.univie.ac.at/cgi-bin/RNAWebSuite/RNAfold.cgi). RNAfold is a predictor of the secondary structure and indicates the thermodynamic characteristics of each molecule, such as Minimum Free Energy (MFE), diversity, and frequency of sequences (Hofacker, 2003). The alignments among the hop miRNA precursors and their homologs were also performed using RNAalifold (ViennaRNA Services).

### 2.3. *In silico* analysis of miR156 and miR172 targets

In order to infer which members of miR156 and 172 families predicted in the hop genome are targeting SPL and AP2 transcripts, we used the psRNATarget tool (https://www.zhaolab.org/psRNATarget/) (Dai et al., 2018; Dai and Zhao, 2011). The mature sequences were used as queries against candidate transcript targets retrieved from Hopbase (http://hopbase.cgrb.oregonstate.edu/) (Padgitt-Cobb et al., 2021). It was considered those microRNAs that presented a Maximum expectation (Exp) value between 0 and 1. In addition, to confirm the presence of SPL and AP2 or AP2-like domains in the putative target transcripts, their protein sequences of each were inspected for conserved domain by using the NCBI conserved domain database online tool (https://www.ncbi.nlm.nih.gov/Structure/cdd/wrpsb.cgi).

### 2.4. Identification PEBP family members, selection of FT-like genes in the hop genome and cis-regulatory elements analysis

We used the BLAST v.2.11.0 (Altschul et al., 1997) to scan the hop proteome searching for PEBP proteins sequences. The FT protein from *A. thaliana* was used as a BlastP query against putative proteins obtained previously by our group (Márquez Gutiérrez et al., 2022). In parallel, BlastP was also carried out against the hop proteome retrieved from the HopBase platform. We only considered PEBP proteins, those sequences presenting a conserved domain verified by domain inference analyses. Redundant proteins reported in the same locus were collapsed after manually curating genomic loci with IGV (Thorvaldsdóttir et al., 2013).

PEBP protein sequences from multiple species retrieved from the NCBI database, along with those from *H. lupulus* described above, were used in our phylogenetic analysis. The multiple sequence alignment (MSA) was performed with MAFFT v.7.475 (Katoh et al., 2002). The MSA was evaluated with GUIDANCE 2 v.2.02 (Sela et al., 2015) to remove low confidence sequences. Both steps were run under default parameters. Phylogenetic trees were inferred with PHYLIP v.3.696 (Felsenstein, 1989), using the Jones-Taylor-Thornton matrix and neighbor-joining clustering method (Saitou and Nei, 1987) over 5,000 bootstrap replicates. Finally, the phylogenetic tree was visualized with the online tool interactive Tree Of Life (iTOL) (Letunic and Bork, 2021) and the hop PEBP proteins were classified into subgroups according to the Arabidopsis PEBP protein classification (Jin et al., 2021).

The PEBP hop proteins that clustered in the FT-like clade were further analyzed to determine if they are a potential floral inducer or not. First, the global alignment with orthologous sequences was visualized in R v.4.3.0 using the “ggmsa package”. This was important to determine the presence of amino acids in critical position which has been determined to confer FT floral inducing activity (Ahn et al., 2006; Ho and Weigel, 2014; Jin et al., 2021). Finally, cis-regulatory elements of the FT-like loci were scanned up to 6,000 nt upstream from the transcription start site (tss) with the PlantCARE database (Lescot et al., 2002).

### 2.5. RNA extraction and Real-Time Quantitative PCR (RT-qPCR) analyzes

Total RNA from leaves were extracted according to de Oliveira et al. (2015). To eliminate DNA contamination, RNA samples (5 μg) were treated with DNase I using the Turbo DNA-free Kit (Ambion). RNA integrity was analyzed in 1 % agarose gel, and RNA content, as well as purity, were accessed by spectroscopy in which all samples showed 1.8<OD260/280<2.2 (NanoVue GE Healthcare, Munich, Germany).

The stem-loop method (Varkonyi-Gasic et al., 2007) was used for cDNA synthesis, following the procedures by Cardoso et al., (2018) with some modifications: using the ImProm-IITM Reverse Transcriptase (Promega), in which 1 μg of treated RNA was adjusted in a volume of 8.5 µl with RNAse-free water. To the treated RNA was added 1.25 µl of oligo-dT primer, 2.5 µl of each specific stem-loop RT primer (10 µM) and 1.25 µl of the dNTP (10 µM). To perform the qPCR of FT-like genes, 1 μg of the total RNA was reverse transcribed into cDNA using the High-Capacity cDNA Reverse Transcription Kit (Thermo Fisher Scientific, Waltham, MA, United States), according to the manufacturer’s protocol. Relative gene expression was calculated based on the 2-ΔΔCT method (Livak and Schmittgen, 2001) and normalized against *S-ADENOSYLMETHIONINE SYNTHETASE* (*HlSAMS*-000185F.g19.t1) and *TRANSLATION ELONGATION FACTOR 1-ALPHA* (*HlTEF1A*-000060F.g21.t1). These reference genes were selected based on their expression stabilities across public RNAseq libraries and all the primers used are available in Table S1.

Real-Time Quantitative PCR (RT-qPCR) was performed with the Rotor-Gene SYBR® Green PCR Kit (Qiagen), on a Rotor Gene-Q(R) thermocycler (Venlo, Netherlands) using three biological repetitions runned in three technical replicates as described by López et al., (2022). The expression rate and the confidence intervals were calculated according to the method proposed by Steibel et al., (2009), which considers the linear mixed-effect model. Residuals were verified to be normally distributed and the graphics were drawn with R v.4.3.0.

## 3. Results

### 3.1. Phenological analysis during the hop growth

The hop phenological cycle under a subtropical climate accompanied along one year is shown in Figure 1. We found that hop presented a different growth cycle in subtropical climate compared to temperate climates (Roßbauer et al., 1995). In one year it was observed four complete cycles of hop growth (Figure 1A) starting with the rhizome buds break (Figure 1B), the formation of a new bine lasts for one week (beginning of shoot-growth, Figure 1C). Then, the growth was linear until plants found support and started to grow in the clockwise direction (Figure 1D). Although very rare, some plants independently of cultivar started to grow secondary branches (from 18 or 22th node) when 25 or 30 nodes were developed.

**Figure 1.**
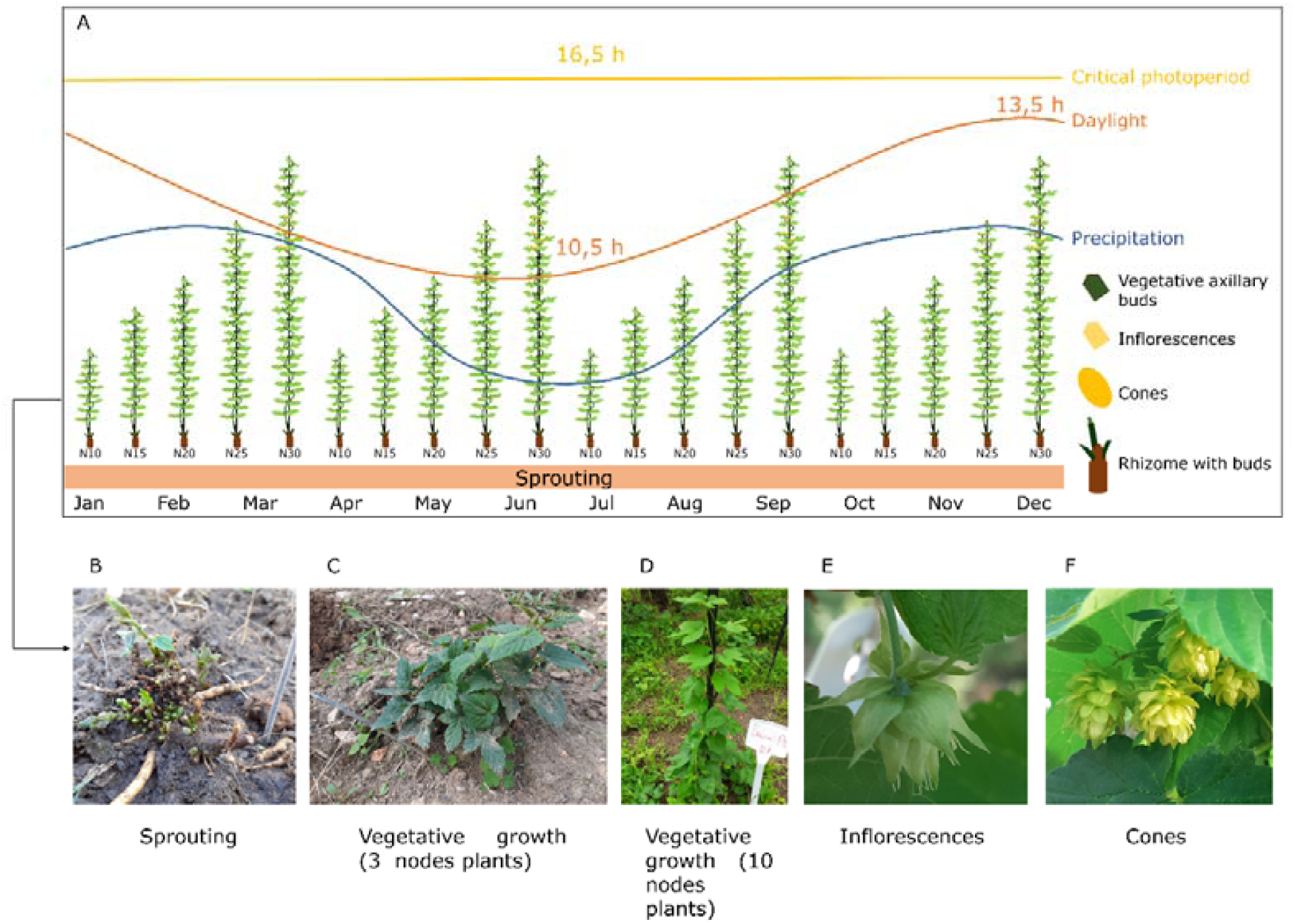
Developmental pattern of hop in subtropical Brazilian conditions. A) The photoperiod (orange line) increased from a minimum of 10.5 hours on 10th June to a maximum of 13.5 hours on 21th December, which is always below the critical photoperiod of 16.5h (yellow line). The rainy period starts in September and extends to May being followed by the dry season (blue line). The phases are shown as follows: B) sprouting, C) vegetative growth in three nodes stage, D) vegetative growth with ten nodes, E) inflorescence stage and F) cones stage.

The plants began to flower when 25-30 visible nodes were formed (Figure 1E). The flowering was observed in any time of the year, being dependent on plant size, in terms of number of nodes, and genotype. For example, plants of the NB cultivar started to flower when 25 nodes were developed and flowering was only verified in plants with 30 nodes for Chinook and Millenium genotypes. The first inflorescences emerged in the 18 to 22^th^ nodes in the axillary buds of primary bines (Figure 1E). The plants with secondary branches also presented flowers in their axillary buds. Interestingly, the flowering stopped when plants reached 30 or 35 nodes and the last inflorescences emerged in the nodes 28 or 30^th^, although the bines continued to grow until the plants started to senescence.

The inflorescence stage lasts 1-2 weeks (Figure 1E), ending when the stigmas begin to degrade. After that, the cones began to develop and lasted for one month (Figure 1F). Then, it started to degrade and fall while the plants were senescing. When the aerial parts were removed, new buds from rhizomes developed into new bines and the cycle started once again at any point of the year (Figure 1). In addition, it is important to highlight that during the winter (from June-August), which is cold and dry in this region, the plants were irrigated, so the deficit of water in hop development was not evaluated here.

Another aspect of hop development is that this plant presents heteroblasty (Fric et al., 1991), which could be observed in the leaves of the cultivar NB during the vegetative development (Figure 2). It is possible to see a heart shaped juvenile leaf leaf (Figure 2A) becoming trilobed (Figure 2B), which occurs usually, from the 7th or 8th nodes (Figure 2C and D) and marks the transition to the mature stage. We found that when NB plants, growing in the evaluated conditions, reached the 25 node stage they began developing the inflorescences in the axillary of the principal branch (Figure 2E, F).

**Figure 2.**
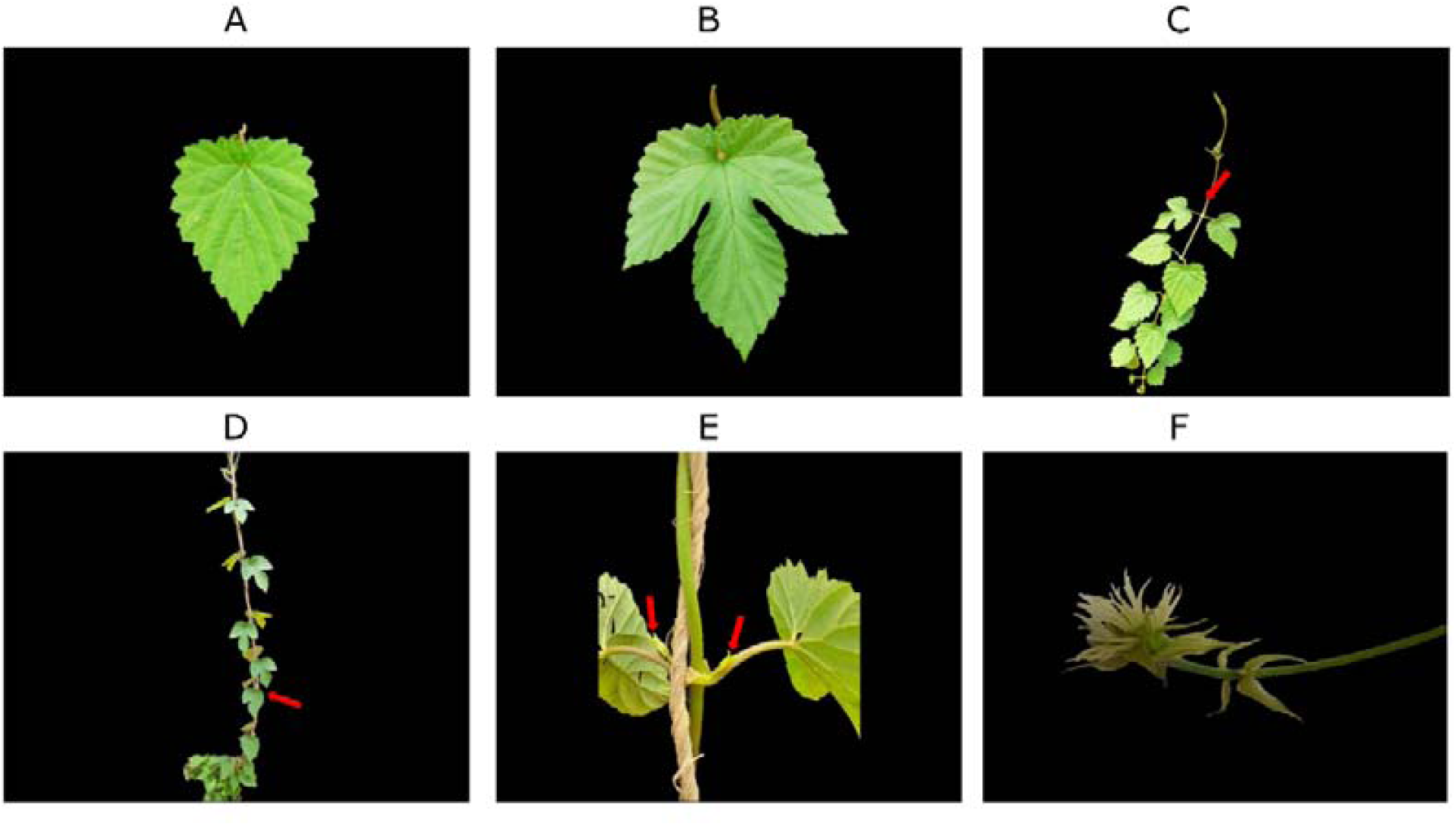
Heteroblasty in hop leaves observed for the NB genotype during the vegetative and reproductive development. Heart shaped juvenile leaf (A). Trilobed leaf (B). Change from heart shaped juvenile leaf to trilobed leaves in 7th node (red arrow) (C). Change of heart shaped to trilobed leaves from the 7th node to the apical meristem (red arrow) (D). Inflorescences in the 20th axillary node (E). Inflorescences in the 20th axillary node one week after blooming (F).

### 3.2. Identification of precursors and mature miRNAs

A total of 137 precursors of miRNAs and 188 mature miRNAs were identified representing 75 conserved families of MIRNAs (Table S1), 35 more than the 40 family previously found (Mishra et al., 2016). Most families contain only one mature sequence being the miR156, the family with the highest number, fifteen, followed by miR171 and miR169 with fourteen and eleven mature sequences, respectively (Table S1). Furthermore, the sequence length ranged from 18 to 24 nucleotides (nt) and found in common had more than 80% identity with the main homologues.

### 3.3. Precursors and mature miRNAs 156 and 172 in hop and prediction of their potential targets

To explore in detail the microRNAs related to reproductive development, we selected the identified members of the microRNAs 156 and 172 families in the hop genome (Table S1). We found eight members of the MIR156 family that codified precursors, with length ranging between 96 and 263 nt (Table S1), three of them were found to produce mature copies of hlu-miR156d. A total of fifteen mature sequences were predicted to have originated from the pre-miR156 precursors with sizes that vary from 18 to 22 nt (Table S1). The precursor hlu-miR156a only originates one mature sequence in its 3’ arm. For the MIR172 family, four members were found codifying precursors with sizes between 177 and 253 nt, from these, the member hlu-miR172b is duplicated in the genome (Table S1). These members of the MIR172 family give place to eight mature miRNAs (3p and 5p) that vary in size between 19 and 23 nt (Table S1).

Characterization of these two highlighted families was performed to compare the identified miRNAs with the corresponding homologues (Figure S1 and S2). The secondary structures were shown to form hairpins. Regarding the alignments, the sequences showed more than 80% identity in the most important part of the miRNAs, the mature sequences, which is the part that effectively binds to mRNA targets in plants.

The prediction of targets of mature miRNAs156 and 172 in the hop genome revealed that only hlu-miR156a-3p presented a 100% of match (or expectation of “0”) with six SPL genes of hop (Table S1). These results are in agreement with targets described in other plants (Wu et al., 2009). Regarding to the eight miRNA172, two out of them, hlu-miR172-3p and hlu-miR172d-3p, presented an expectation of 0.5 and 1 with four AP2 and three AP2-like of hop (Table S1), which means that only one or two nucleotides does not match with targeting regions in these genes. In this case, hlu-miR172d-3p targeted three AP2-like genes. In addition, hlu-miR156a-3p, hlu-miR172-3p and hlu-miR172d-3p appear to regulate these target genes through cleavage. According to these results, these microRNAs were selected to have their expression evaluated during hop development.

### 3.4. RT-qPCR of miR156 and miR172 during leaf development

The expression of three miRNAs (hlu-miR156a-3p, hlu-miR172d-3p and hlu-miR172-3p) was accessed by RT-qPCR in leaves with different ages and node position of three hop cultivars (Figure 3). Comparing leaves at different nodes, the hlu-miR156a-3p expression decreases significantly from the 10 nodes plants to the 25 nodes plants for all the three genotypes (Figure 3A). Contrary to this pattern, the expression of hlu-miR172d-3p and hlu-miR172-3p increases significantly from 10 node plants to 25 node plants, especially for the Chinook and Millenium genotypes, whereas this increasing could not be verified for the NB (Figure 3B and C). In this genotype, hlu-miR172d-3p showed significant expression only in plants with 20 nodes when compared to plants with 10 nodes (Figure 3B). Nevertheless, the expression of hlu-miR172d-3p did not exhibit statistical differences among different plant sizes for the cultivar NB (Figure 3C).

**Figure 3.**
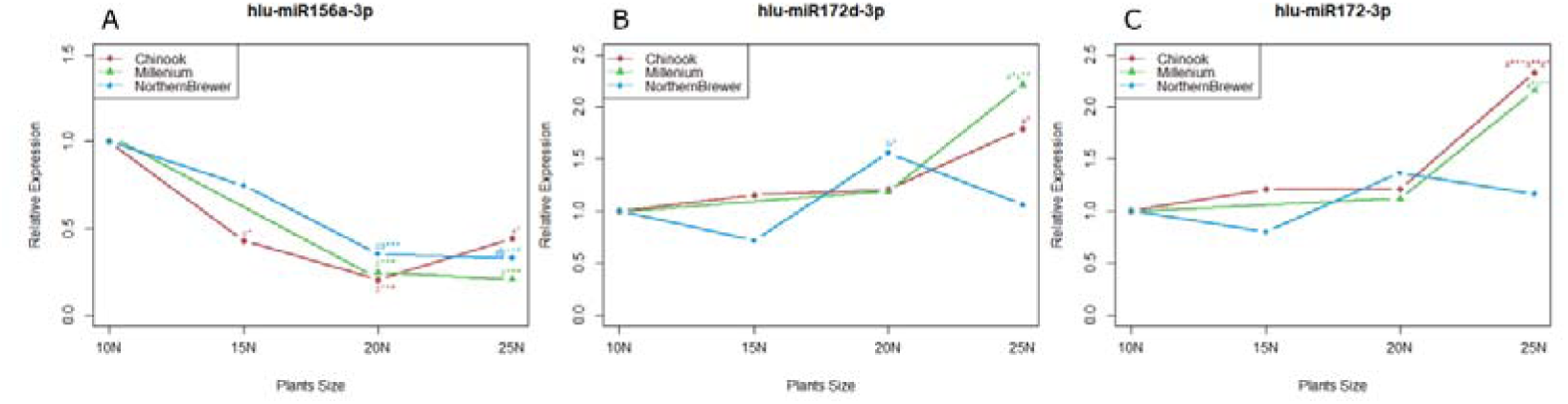
Relative expression of hlu-miR156a-3p (A), hlu-miR172d-3p (B) and hlu-miR172-3p (C) in three hop genotypes: Chinook, Millenium and NB in leaves from plants with 10, 15, 20 and 25 nodes. The letter “a” means significant differences compared with 10 nodes plants, the letter “b” means significant differences compared with 15 nodes plants, and the letter “c” means significant differences compared with 20 nodes plants. The asterisks indicate the significance level p=0 ‘***’ p=0.001 ‘**’ p=0.01 ‘*’.

### 3.5. Expression analyses of FT-like genes and its promoter cis-regulatory elements

To determine the expression profile of *FT-like* in hop along development and to test the hypothesis that it induced in an age-dependent way, we firstly identified and classified the PEBP proteins. Throughout phylogenetic analysis, we found sixteen PEBP protein (Figure 4A): three in the MFT clade (HlMFT1-3), six in the TFL clade (HlTFL1-6), and seven in the FT-like clade (HlFT1-7). In addition, a sub-clade within the TFL clade that seems to be specific to the Cannabacea family with six members, however members of the BFT subclade were not found in the hop genome (Figure 4A).

**Figure 4.**
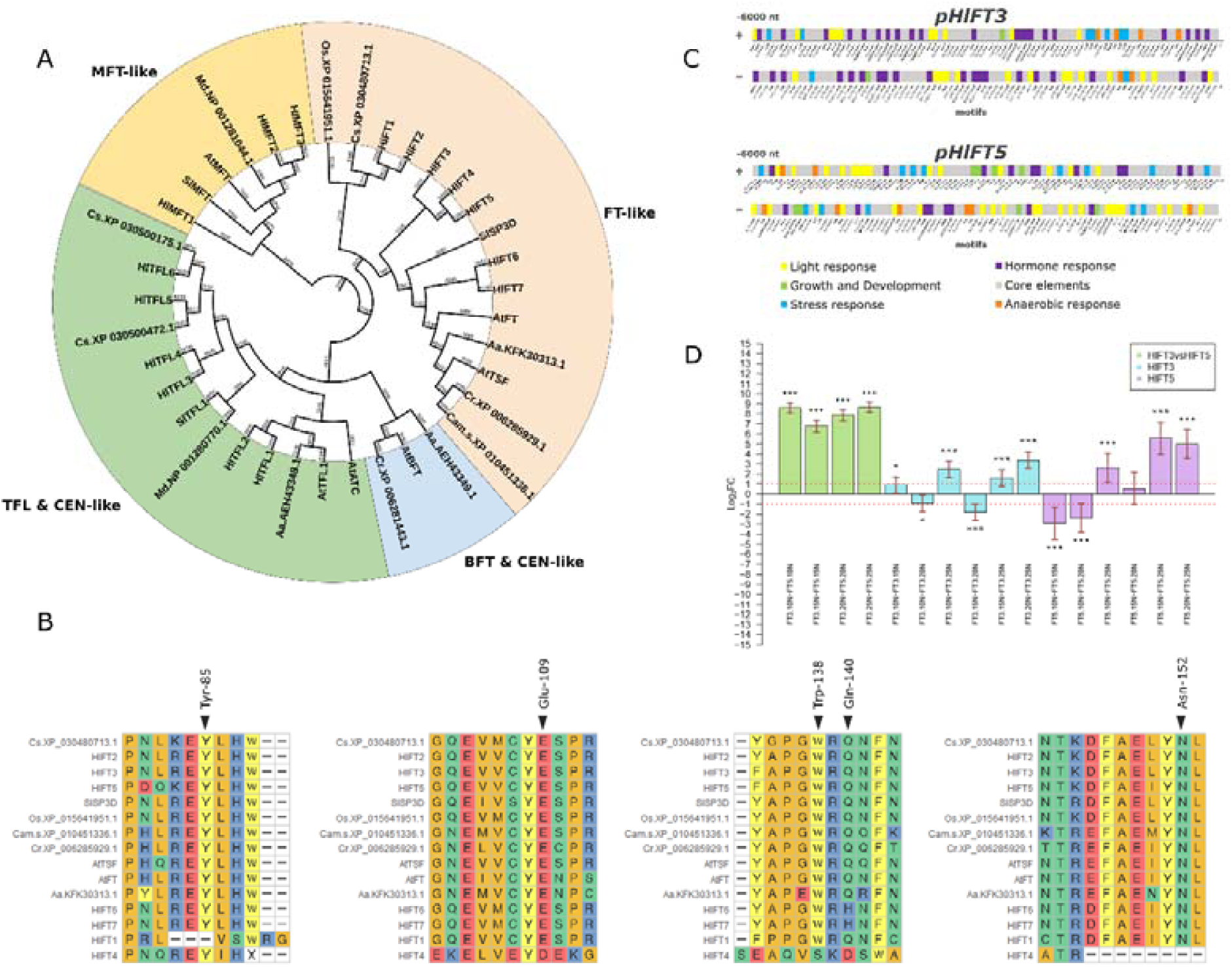
Phylogenetic analysis of PEBP proteins of hop (Hl), *A. thaliana* (At), *Arabis alpina* (Aa), *Capsella rubella* (Cr), *Camelina sativa* (Cam.s), Cannabis sativa (Cs), *Malus domestica* (Md), *Solanum lycopersicum* (Sl) and *Oryza sativa* (Os) (A). Multiple sequence alignment of PEBP proteins that clustered into FT-like clade; the essential amino acids and positions that confer FT activity are shown (B). Cis-regulatory elements predicted in positive and negative strands of 6000 nt upstream of translation start site of *HlFT3* and *HlFT5* (C). RT-qPCR expression of *HlFT3* and *HlFT5* during NB cultivar development. The asterisks indicate the significance level p=0 ‘***’; p=0.001 ‘**’; and 0.01 ‘*’. Bar plots represent the relative difference in expression in log2 of fold change (Log2FC) in relation to the first term described. Green bars represent comparisons between both genes (*HlFT3* vs *HlFT5*) in each stage analyzed, for example *HlFT3* (first term of comparison) is always more expressed than *HlFT5* (second term). Each bar (blue and lilac) represents one comparison in gene expression between plant size (number of nodes). Error bars in red are the 95% confidence interval calculated using the LMM methodology, a gene is deemed Differentially Expressed (DE) if its confidence interval does not cross the number 0 in the y axis (D).

The presence of certain conserved amino acids at specific positions of FT-like proteins (Try-85, Glu-109, Trp-138, Gln-140 and Asn-152) is necessary to its function as floral activators (Ahn et al., 2006; Ho and Weigel, 2014). Thus, we analyzed these important regions of conserved amino acids and found that there are two PEBP proteins with all the required positions filled with the specific floral inducing components (potential florigens). These proteins were named HlFT3 and HlFT5 with 176 and 175 amino acids, respectively (Figure 4B).

Arabidopsis *FT* is controlled by the circadian rhythm (Adrian et al., 2010), thus in order to explore related motifs in the *HlFT3* and *HlFT5* promoter we checked the cis-regulatory elements in 6000 nt upstream of the translational start sites. Results showed that both genes could respond differently to environmental changes, since the promoter of *HlFT3* contains more motifs to hormone response whereas *HlFT5* contains more motifs to light response (Figure 4C).

Considering both strands of the 6,000 nt upstream region of *HlFT3* we identified thirty nine hormone responsive motifs, twenty four motifs that respond to light, ten to stress response, six to anaerobic response and two growth and development motifs (Figure 4C). The most represented hormone responsive motifs were the ones related to jasmonic acid methyl ester (MeJA) TGACG-motif (ten occurrences) and CGTCA-motif (ten). Other hormone response motifs such as the ethylene responsive motifs, represented by the ERE (eleven), the ABRE abscisic acid response motifs (four), the salicylic acid responsive TCA-element (two), auxin response by TGA-element (one) and the P-box gibberellin response motif (one) were also found within the *HlFT3* cis-regulatory elements. The light responsive elements in the *HlFT3* promoter were represented by the Box-4 (seven), AE-box (four), G-box and GT1-motif (three each), TCCC and TCT motifs (two each), and Sp1, GTGGC, chs-CMA1a, and AT1 motifs (one each). In the case of stress response elements, the most represented was the LTR (seven), a cis-acting element involved in low-temperature responsiveness.

In contrast to the promoter region of the *HlFT3*, the 6,000 nt upstream of HlFT5 is preferentially enriched for light responsive elements instead of hormone responsive. The most abundant light responsive elements were the Box-4 motif (seven occurrences), followed by the motifs TCT and GATA (four each), the elements I-box, E-box, GT1, and G-box (three times each), the motifs GA, GATA, and MRE were distributed (two times each), while the motifs ATC, GARA, LAMP and LS7 had a single occurrence. Regarding the hormone responsive elements, the most abundant in the *HlFT5* promoter was related to abscisic acid responsiveness, ABRE (five), followed by the ERE motif (four), the TGACG-motif (two) and CGTCA-motif (two), whereas a gibberellin-responsive element (P-box, two occurrences) was identified, and an element responsive to auxin was also found (TGA-element, two occurrences). The others motifs identified in the *HlFT5* promoter were these related to anaerobic response (ARE, seven occurrences), those that respond to stress like WUN-motif (six), the TC-rich repeats motif (three), LTR (two), and MBS (one). Finally, regarding the plant growth and development elements, it was possible to identify the motif CAT (three), the CCGTCC-box motif (one), the plant AP-2-like motif (one), the AC-I motif (one), and the AC-II motif (one).

In order to explore the FT-regulatory pathways along hop development and correlate it with miR156 and miR172 expressions (Figure 3), we analyzed the expression of *HlFT3* and *HlFT5* in leaves of plants with different sizes (10, 15, 20 and 25 nodes). In figure 4D it is observed that in all stages analyzed, *HlFT3* is more expressed than *HlFT5* for almost 250 times (green bars). Comparing the expressions between plant with different node number (blue bars), the expression of *HlFT3* is significantly higher (2 times, p < 0.05) in plants with 10 nodes (10N) than in plants with 15 nodes. In plants with 20 nodes, the expression of *HlFT3* is significantly higher than in plants with 10 nodes (2 times, p < 0.05), and also higher than in plants with 15 nodes (3.5 times, p < 0.001). In all cases, the expression of *HlFT3* is lower in plants with 25 nodes (p < 0.001), indicating that the peak of expression occurs in plants with 20 nodes.

Meanwhile, although less expressed than *HlFT3* in all considering conditions, *HlFT5* is more expressed in 15 nodes plants (8 times, p < 0,001) and in 20 nodes plants (5 times, p < 0,001) than in plants with 10 nodes (Fig. 4D, lilac bars). The expression of *HlFT5* is not different between 15 and 20 nodes plants. Similar to the *HlFT3* expression pattern, in plants with 25 nodes the expression of *HlFT5* decreased (Figure 4D).

The high expression of both *HlFT3* and *HlFT5* in the 20 nodes plants coincides with the low expression of the miR156 and increase of miR172 (Figure 3) together with the exact time of the observed floral transition in Brazilian conditions for NB plants (Figure 1). From that, a model was depicted explaining the floral development and the related miRNA156/172-FT pathway (Figure 5).

**Figure 5.**
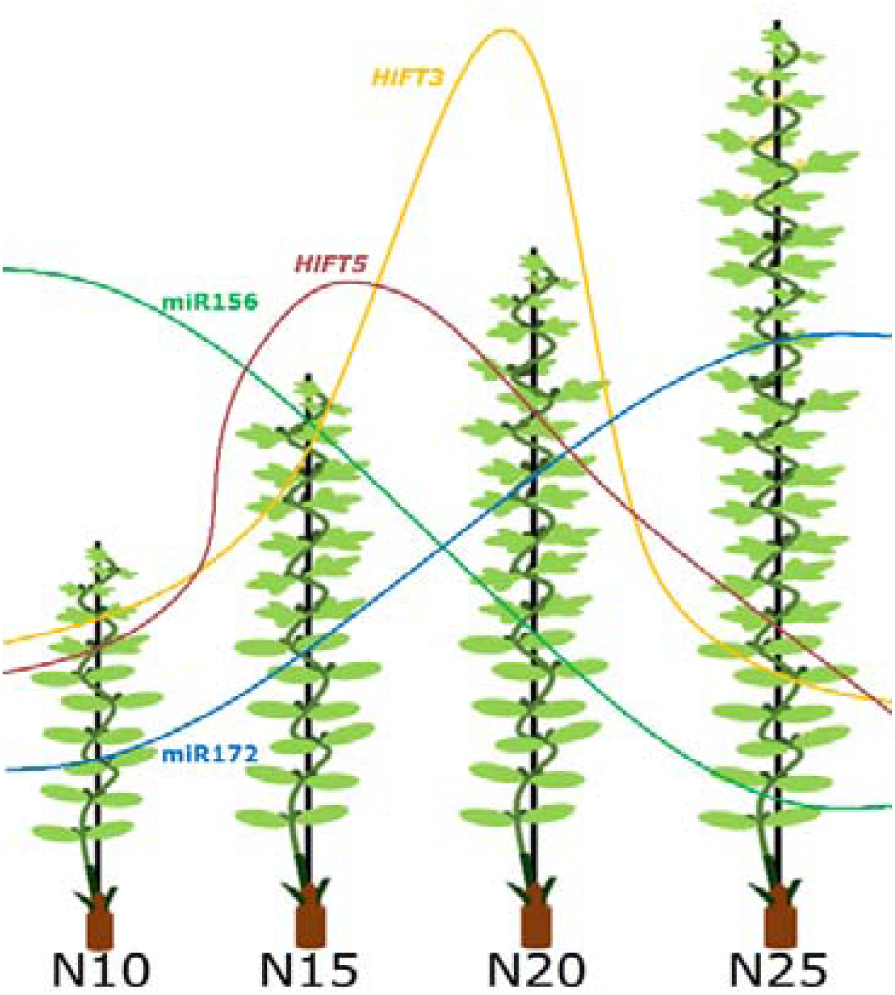
A model of the miRNA156/172 and FT expression explaining the hops floral transition. The high expression of both *HlFT3* and *HlFT5* occurs in the 20 nodes plants (Figure 4D) which coincides with the low expression of the miR156 and increase of miR172 (Figure 3) together with the exact time of the observed floral transition in Brazilian conditions for NB plants (Figure 1A).

Moreover, the differences of expression between *HlFT3* (more expressed) and *HlFT5* suggest that in the evaluated conditions, the *HlFT3* acts as the true florigen. Many hypotheses can be raised for the *HlFT5* homolog function, which could have retained redundant functions to *HlFT3*, sub functionalized or acquired a different function, or be stimulated by different environmental cues, or even lost the function as a florigen. Supporting some of these hypotheses, the different expression pattern of *HlFT3* and *HlFT5* may be a consequence of the contrasting distribution of regulatory elements found in their promoters (Figure 4C) that suggests that *HlFT3* is primarily responsive to hormones whereas *HlFT5* is primarily responsive to light. Thus, further analysis at different growth conditions is necessary to respond to these questions.

## 4. Discussion

Studies of the phenological cycles and phenotypical changes can be accompanied by molecular analysis to better elucidate the intricate routes that dictate the stage transition between species. It is also important for the understanding of how crops have evolved and adapted to different regions in response to particular environmental changes. In other words, how crops perceive environmental signals, such as temperature and photoperiod, to trigger the flowering development and achieve reproductive success. Therefore, interpreting these interconnected abiotic and biotic processes allows us to predict the behavior of a plant in some conditions, and above all, direct breeding programs towards crop production in different regions, as for hop in Brazilian subtropical conditions.

In this work, we combined phenologic, phenotypic and molecular biology analyses to better understand hop development in subtropical conditions. To do so, we first discriminated the different stages of hop development (Figure 1A) and evaluated the interplay between expression patterns of one miR156, two miR172 and two putative *FT-like* (*HlFT3* and *HlFT5*) in four developmental stages of three hop genotypes cultivated in Minas Gerais/Brazil (Figure 1B). Usually, the start of hop growth in temperate regions begins with the shoot development from buds in the rhizome after the winter period known as endormancy or winter arrest (Roßbauer et al., 1995). In the conditions studied here, when the rhizome buds reach their maximum size, they enter a stage of inactivation that ends with the removal of the aerial parts, triggering the development of new aerial parts (Figure 1C). This phenomenon corresponds to a state of paradormancy or dormancy by correlation, in which the buds are inhibited by the presence of other organs (Falavigna et al., 2019). Moreover, in the North hemisphere the hop floral development starts 90 days after the sprouting and the flowering occurs at the end of summer (Fric et al., 1991; Roßbauer et al., 1995). Contrasting to this hop developing pattern, in the subtropical conditions, we verified that vegetative growth occurs at any time of year and flowering starts once the plants achieve a specific size (more than 25 nodes) in a genotype-dependent manner (Figure 1A).

The flowering of hops is induced when day length is shortened from the 16.5 h (critical point) and plants are competent to perceive this stimulus, once a minimum number of nodes are formed (Thomas and Schwabe, 1969). Accordingly, we observed that the hops cultivated in Brazil are able to flower at any time of the year, but it still depends on plant size or maturity (Figure 1). Moreover, it has been reported that flowering time is genotype-dependent. Early flowering genotypes require a smaller number of nodes to perceive the stimuli, as noted by Thomas and Schwabe in 1969. In addition, Bauerle et al. (2019) also reported that all genotypes used in his study are considered ‘ripe to flower’ when there are at least 25 visible nodes. Thus, it is plausible to assume that in Brazil the photoperiod is always inductive, since day length is always below the critical point, and our phenological observations suggest that the control of hop flowering in subtropical regions mostly depends on endogenous cues, as developmental stage.

Previous reports of hops cultivated in Florida, USA (a subtropical region) observed that the Cascade genotype bloomed as early as 26 days after sprouting when the plants reached 15 node-stage (Acosta et al., 2021). Furthermore, together with the extended flowering window, the senescence of aerial parts overlapped with the development of younger flowers ultimately causing the unevenness in cone development. It was also noted the absence of a distinct lateral shoot formation stage (Acosta et al., 2021). Accordingly, in this work, we observed a similar behavior of the three studied hop genotypes, which may be due to the continuous maintenance of the inductive photoperiod. These results point to a challenge in producing hop flowers in subtropical regions but also the endogenous factors associated with the flowering control that could direct breeding programs.

The hops age and developmental stage (measured as the number of bine nodes) seems to be the major factor that dictates the floral induction in subtropical regions. The physiological age of a plant can be endogenously perceived by the accumulation dynamics of miR156 and miR172 during plant development (Wang, 2014; Wu et al., 2009). Previously has been reported that the molecular function of these conserved miRNAs is the same between the land plants (Axtell et al., 2007). Therefore, the targets identified here (Table S1) are likely to participate in those developmental phases. In agreement, we observed the decreasing of hlu-miR156a-3p during hop development (Figure 3A). This behavior coincides with an intermediary stage between the transition from juvenile to adult phase where we observed leaf morphology changes from heart shaped to trilobed margins in the 7th or 8 nodes (Figure 2), similarly to *P. edulis* (Silva et al., 2019). In *A. thaliana* it was determined that leaf identity depends on balance between SPL/miR156 in the SAM (Fouracre and Poethig, 2019). These investigators also demonstrated that the decrease of SAM size is due to an increase of SPL transcription factors, making the SAM larger in response to a higher level of miR156. Previously, reduction of SAM size was observed in female and male hop plants during development (Shephard et al., 2000). Therefore, it is plausible that in hop, hlu-miR156a-3p is associated to this process since their targets are members of the SPL transcription factor family (Table S1).

The down-expression observed for hlu-miR156a-3p also depends on the node’s position, since the analysis was performed on leaves from 5th, 10th, 15th, and 20th nodes, in plants with 10, 15, 20 and 25 nodes, respectively (Figure 3A). This is in agreement with the cell division of SAM in *A. thaliana* which triggers repressing epigenetic marks in the MIR156 locus, showing that the apical parts of a plant are physiologically older than the basal parts (Cheng et al., 2021). This statement is as interesting as counterintuitive, since the apical parts are developed latter (younger) from the SAM. Nevertheless, the logic lies in the number of cell divisions in the SAM the more divisions the more epigenetic marks are added causing additional molecular markers of age in MIR156 locus. Thus, this phenomenon explains the preferential flowering near to the tip in hop plants in contrast to the typical juvenile morphology in the first leaves.

In Arabidopsis it has been demonstrated that the expression of miR156 is promoted by the AP2, which in turn is a miR172 target (Yant et al., 2010), therefore the miR172 acts in opposition to miR156, promoting the floral transition (Teotia and Tang, 2015; Wang, 2014; Wu et al., 2009). In this work, we show *in silico* evidence that hlu-miR172d-3p and hlu-miR172-3p are targeting seven members of AP2 or AP2-like (Table S1). High levels of hlu-miR172d-3p expression was observed when NB plants achieved 25 node stage (Figure 3B), while both mature miRNAs hlu-miR172d-3p and hlu-miR172-3p were significantly expressed in plants with 25 nodes for Chinook and Millenium. This could explain why NB plants flowered when 25 nodes were visible, while both Chinook and Millenium cultivars flowered later, once 30 nodes were formed. Therefore, it is likely that the differences of flowering time between the analyzed genotypes is due to differences of miR172 expression. Perhaps the expression of miR172 in plants with 20-25 nodes is needed to promote the up-regulation of FT in inductive conditions, since their targets in A. thaliana, are AP2 or AP2-like transcription factors known inhibitors of the FT transcription (Hu et al., 2021; Mathieu et al., 2009).

*In silico* analyses revealed sixteen PEBP proteins being seven clustered in the FT-like clade (Figure 4A). From this, we selected HlFT3 and HlFT5, that presented the conserved amino acids tyrosine in the 85th position, glutamate in the 109th, tryptophan in 138th, glutamine in 140th and asparagine in 152th (Figure 4B). Justifying this selection, previous studies have demonstrated that point mutations of some of these amino acids can confer an TFL-repressor activity in FT protein (Ahn et al., 2006; Ho and Weigel, 2014), thus, in hop, these two genes have the potential to act as florigen.

Cis-regulatory elements analysis of HlFT3 and HlFT5 show several motifs related to different environmental conditions in their promoter regions (Figure 4C). We found seven Box-4 and three G-box distributed along 6,000 nt upstream of *HlFT3*. Which could be related to enhancing the FT transcription in response to blue light during the dusk. Meanwhile, the presence of cis-regulatory elements related to Jasmonic acid, ethylene, and ABA in the HlFT3 promoter could be related to the flowering of plants in response to senescence state as previously observed by Acosta et al (2021). Moreover, the presence of cis regulatory elements related to low temperature in both *HlFT3* and *HlFT5* promoters could also be related to cold responses in a photoperiod longer than 16,5 h light (Thomas and Schwabe, 1969). Therefore, future studies must explore the functional redundancy of HlFT3 and HlFT5 in response to different environmental conditions and its impact on the flowering.

The *HlFT3* and *HlFT5* have their expression varying along the hop developmental stages suggesting an induction window in plants between 15 and 20 nodes in the NB genotype (a model is available in Figure 5). The statistical significant up-regulation of the *HlFT3* expression coincides with the higher accumulation of hlu-miR172d-3p/hlu-miR172-3p and lower of hlu-miR156a-3p (Figure 3) agreeing to similar results in other species and pointing the conservation of the regulatory motif miR172/AP2/FT (Castillejo and Pelaz, 2008; Hu et al., 2021; Mathieu et al., 2009). This expression pattern of HlFT3 could explain the emergence of inflorescences in the 20-25th nodes of NB plants and the absence of new inflorescences from the 26th node when both HlFT3 expression abruptly decreases. This indicates that hop plants growing in inductive conditions need to achieve a cultivar dependent minimum size to perceive the short-day stimuli, and once the size is achieved, the *HlFT3* expression is promoted. In agreement, we found a cis regulatory element responsive to AP2-like at 5110 nt upstream of the *HlFT5* transcription start site, which suggest a transcriptional regulation by AP2-like proteins, probably a targeted of both hlu-miR172-3p and hlu-miR172d-3p (Table S1).

## 5. Conclusions

Our work indicates that, while considering the differences in hop phenology depending on the cultivated region as subtropical conditions, the sequential stages of development must be described in terms of phenotypic changes and not chronological or season periods. The main conclusion is that in floral inductive photoperiod, below the 16.5 hours in the tropical zones, the hop flowering depends primarily on endogenous signals. Thus, hop should be considered a short-day plant and able to develop flowers at any time of the year in such conditions. This feature is corroborated by our molecular results, in which, the higher *HlFT3* expression coincides with the down-accumulation of miR156 and up-accumulation of miR172 along development, a regulatory motif largely known to be involved in floral transition and conserved in different species. On the other hand, this also means that in subtropical conditions, hops develop and produce flowers dispersed throughout the year, which may imply lower production. Thus, our work contributes to comprehension of hop development and reproductive transition and opening perspectives of managing practices to its cultivation in tropical and subtropical regions that will be different from traditional methods applied in temperate regions.

## 6. Acknowledgements and funding

The authors thank the Universidade Federal de Lavras (UFLA/Brazil) and members of the Laboratório de Fisiologia Molecular de Plantas (LFMP, UFLA/Brazil) for structural support of the experiments and analysis. The Laboratory of Plant Molecular Physiology (LFMP) is partially supported by the Coordenação de Aperfeiçoamento de Pessoal de Nível Superior (CAPES) and the Conselho Nacional de Desenvolvimento Científico e Tecnológico (CNPq). R.M.G. received a PhD scholarship from the PAEC-OAS-GCUB-Program–Organization of American States and CNPq (grant 149043/2019-8).

## 7. Authors’ contributions

R.R.O., R.M.G., and A.C-J conceptualized the project; R.M.G conducted all data research and analyses; T.H.C.R supported for bioinformatic analyses; R.R.O. and A.C-J supervised experiments and analyses. K.K.P.O supported in the molecular analyses; J.V.N.S., T.C.A., L.R.d.A., M.d.S.G., and M.d.S.G contributed to the miRNAs prediction; R.M.G wrote the first version of the manuscript and R.R.O. contributed to final version writing; all co-authors corrected and contributed to writing of the final manuscript version.

## 8. Declaration of Competing Interest

The authors declare that there are no conflicts of interest.

## 9. Data availability

Main data supporting the findings of this study are available within the paper and within its supplementary materials published online. The raw data used for analyses and figures are available from the corresponding author, Antonio Chalfun-Júnior, upon request.

## Notes

### Competing Interest Statement

The authors have declared no competing interest.

